# Further evidences of an emerging stingless bee-yeast symbiosis

**DOI:** 10.1101/2023.03.08.531834

**Authors:** Gabriela Toninato de Paula, Weilan Gomes da Paixão Melo, Ivan de Castro, Cristiano Menezes, Camila Raquel Paludo, Carlos Augusto Rosa, Mônica Tallarico Pupo

## Abstract

Symbiotic interactions between microorganisms and social insects have been described as crucial for the maintenance of these multitrophic systems, as observed for the stingless bee *Scaptotrigona depilis* and the yeast *Zygosaccharomyces* sp. The larvae of *S. depilis* ingest fungal filaments of *Zygosaccharomyces* sp. to obtain ergosterol, which is the precursor for the biosynthesis of ecdysteroids that modulate insect metamorphosis. In this work we verified that nutritional fungal symbioses also occur in other species of stingless bees. We analyzed brood cell samples from 19 species of stingless bees collected in Brazil. The osmophilic yeast *Zygosaccharomyces* spp. was isolated from eight bee species, namely *Scaptotrigona bipuctata, S. postica, S. tubiba, Tetragona clavipes, Melipona quadrifasciata, M. fasciculata, M. bicolor* and *Partamona helleri*. These yeasts form pseudohyphae and also accumulate ergosterol in lipid droplets, similar to the pattern observed for *S. depilis*. The phylogenetic analyses including various *Zygosaccharomyces* revealed that strains isolated from the brood cells formed a branch separated from the previously described *Zygosaccharomyces* species, suggesting that they are new species of this genus and reinforcing the symbiotic interaction with the host insects.

**Importance:** Benefits exchanged in insect–fungus mutualisms include nutrition, protection, and dispersal. Fungal nutritional roles are well described for some eusocial insects, such as fungus growing ants and termites, but similar interaction in stingless bees was so far observed just in *Scaptotrigona depilis*. Here we expand the knowledge of yeast-bee symbiosis by analyzing the presence, cell morphologies, lipid accumulation and phylogenetic relationships of fungi isolated from brood cells and other locations of bee colonies. *Zygosaccharomyces* isolates were recovered from 42% of the bee species assessed, and probably represent new species showing pseudohyphae formation and lipid accumulation similar to *S. depilis* associated *Zygosaccharomyces* strains. The phylogenetic analyses suggested an evolutionary adaptation of *Zygosaccharomyces* spp. to the brood cell environment to provide nutritional benefits for the developing insect. Stingless bees play important ecosystem services, and our results raise the concern that fungicidal agents used in agriculture could disrupt this symbiosis, impacting bee health.

---

Eusocial insects establish several symbiotic interactions with microorganisms, ranging from obligate mutualisms to specialized parasitism (1, 2). The nutritional benefits provided by fungal mutualists is well described for fungus-growing ants, native to the Neotropics, and termites from Africa and Asia (3). Ants of the subtribe Attina cultivate fungi of the families Agaricaceae and Pterulaceae for food (4). Similarly, termites of the subfamily Macrotermitinae cultivate fungi of the genus *Termitomyces*, which are not only the main food source but also provide digestive services (5). Other microbial symbionts also play different roles in such multitrophic interactions, like production of chemical defenses against entomopathogens (1, 2).

Stingless bees (SBs) (Apidae: Apinae: Meliponini) are a monophyletic group of eusocial insects that belong to a larger group of corbiculate bees (6). Meliponini have pantropical distribution and interact with various microorganisms such as bacteria, yeasts, filamentous fungi, and viruses. Indeed, microbial fermentation contributes to important physicochemical characteristics and to the preservation of pollen and honey (7). Interestingly, the benefits provided by a yeast of the genus *Zygosaccharomyces* for the host *Scaptotrigona depilis* has been reported as the first example of nutritional symbiosis in SBs. This osmophilic yeast grows inside the brood cells of *S. depilis* and is eaten by the larvae, an essential process for larval metamorphosis. The fungus accumulates ergosterol, which is used by the developing insect as a precursor for ecdysteroid biosynthesis leading to the proper pupation (8, 9). However, this remarkable interaction had been observed just for one bee species so far, lacking generality among Meliponini (10).

This unprecedented finding of a yeast important for the larval development raised the hypothesis that similar symbiosis might be more spread among SBs. Here, we describe the occurrence of *Zygosaccharomyces* isolates in the brood cells of different SBs species, as well as the phylogenetic relationships and morphological characteristics of these yeasts in comparison with other species isolated from other locations in the colonies of SBs.

Different nest sites of 19 species of SBs, distributed among 12 genera, were assessed for isolation of fungi, resulting in 32 fungal isolates. Twenty-seven were obtained from brood cells, while four were isolated from honey and one from cerumen. All isolates had the 26S, 18S rRNA gene and ITS regions sequenced, resulting in 23 isolates belonging to the genus *Zygosaccharomyces*, seven were identified as *Monascus* spp., one as *Xerochrysium* sp. and one as *Leiothecium* sp. (Supplementary Material - Table S1). Fungal filaments were observed and isolated from the brood cells of eight species of SBs distributed in four genera and eight species (*Scaptotrigona bipuctata, S. postica, S. tubiba, Tetragona clavipes, Melipona quadrifasciata, M. fasciculata, M. bicolor*, and *Partamona helleri*). All collected fungal filaments were identified as belonging to the genus *Zygosaccharomyces*, representing 42% incidence rate of *Zygosaccharomyces* sp. in brood cells of sampled SBs species. *Zygosaccharomyces* were also isolated from honey of four species and from cerumen of one species (Table S1), while fungal filaments were not observed in the brood cells of 10 SBs species. Samples with low amounts of brood cells and the different larval stages could be limiting factors for the observation of *Zygosaccharomyces*, in addition to the difficulty of culturing this microorganism in less complex culture media. Fungal filaments were observed in SBs with disk-shaped brood cells structure (Figure S1A and S1C) and a larger diameter, but were not observed in other species with smaller diameters (Figure S1B, S1D and S1E) and cluster-shaped brood cells (Figure S1D and S1E). The size and shape of the brood cells may influence the visibility of the symbiont fungus.

Cell morphologies of *Zygosaccharomyces* spp. varied according to the site of isolation. Isolates from brood cells form pseudohyphae in glucose-rich culture medium (30 G), while the others have spherical and ovoid cells under the same growth conditions (Figure S2). Pseudohyphae formation can be triggered by low nitrogen levels and is a form of foraging (11). The location of these *Zygosaccharomyces* strains may be a determining factor in the expression of pseudohyphae. The brood cell is composed of cerumen and larval food, and their composition may vary in nature. Cerumen consists only of waxes and resins (12), while larval food consists of water (40-60%), sugars (5-12%), and free amino acids (0.2-1.3%) (13). These water-soluble components make up to 70% of the larval food composition. In addition, pollen and lipids are also present (13). This specific environment with high sugar concentration and low nitrogen content may require some adaptation of the microorganisms, such as the formation of pseudohyphae. It is also hypothesized that pseudohyphae are more easily ingested by the larvae since they are present on the surface of the larval food and have long filaments. This particular morphology of these yeasts (Figure 1) could favor the availability and uptake of nutrients for the larval development.

**Figure 1.**
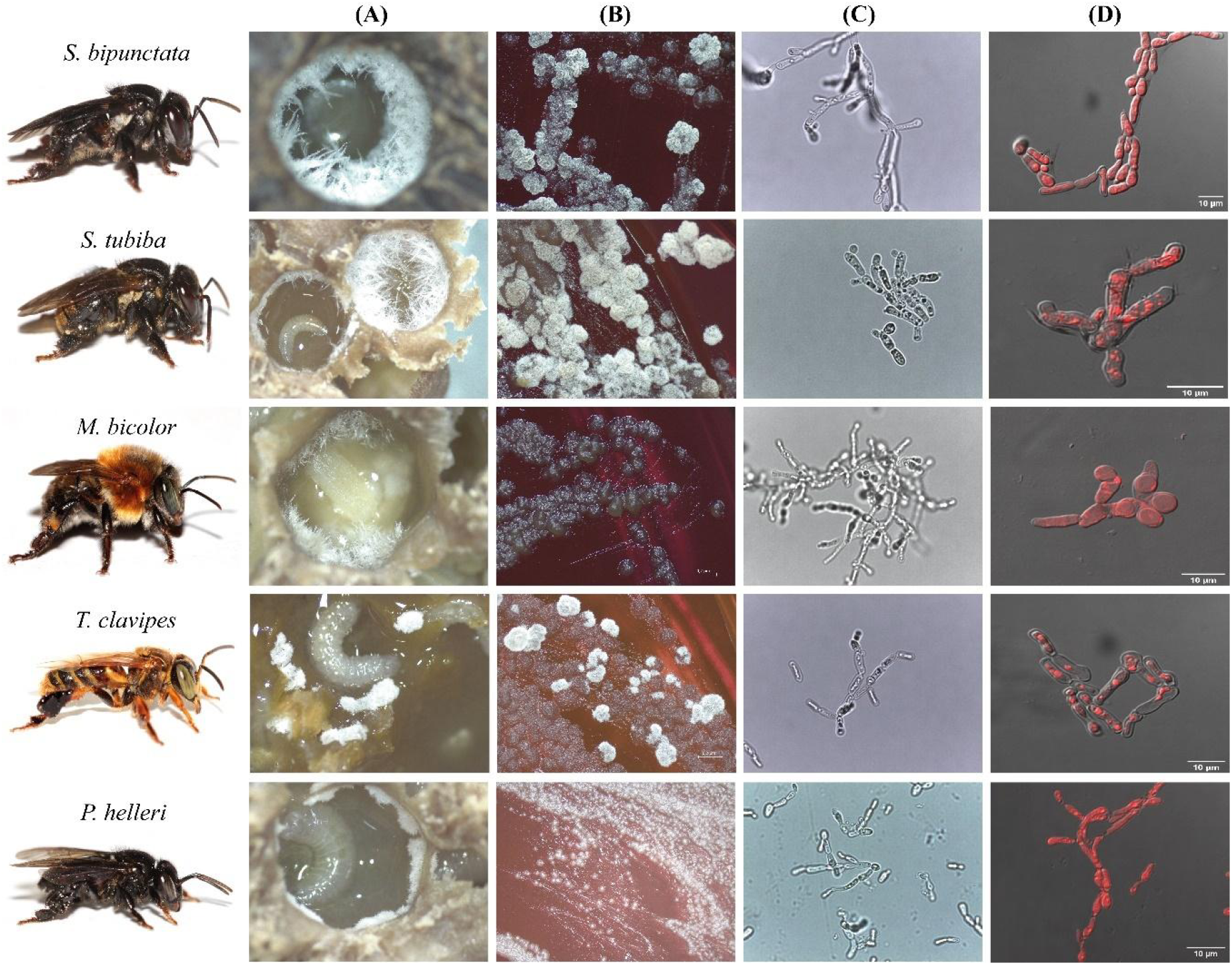
Images of *Zygosaccharomyces* spp. isolates from stingless bees *S. bipunctata, S. tubiba*, *M. bicolor*, *T. clavipes* and *P. helleri*. (A) Fungal filaments present in the brood cells. (B) Microbial growth in Petri dish (14 days) (C) Optical microscopy images (100x). (D) Fluorescence microscopy images. Stingless bees’ photography: credit Cristiano Menezes.

*Zygosaccharomyces* strains from brood cells of different bees showed accumulation of intracellular lipids, as visualized by fluorescence microscopy (Figure 1D). GC-MS analyses of cell extracts of *Zygosaccharomyces* spp. confirmed the presence of ergosterol in all the samples (Figure S3), as previously described for *Zygosaccharomyces* sp. isolated from *S. depilis* (9), reinforcing the nutritional function of the yeasts for SBs larvae.

*Zygosaccharomyces* strains from the brood cells had the lowest query coverage and identity percentages with species from the Genbank database and may represent new species. Phylogenetic tree from Bayesian analyses using the sequences of the D1/D2 domain of the 26S gene was performed on *Zygosaccharomyces* spp. (Figure 2). A combined phylogeny was also constructed using a dataset of the 26S and 18S genes (Figure S4). Phylogenies show a branch consisting of *Zygosaccharomyces* spp. isolated exclusively from the brood cells of different species of SBs. The other lineages of *Zygosaccharomyces* isolated from different nest sites are phylogenetically distant. The branch consists only of *Zygosaccharomyces* spp. strains isolated from the brood cells contain *Zygosaccharomyces* sp. associated with *S. depilis* (9), and is divided into several groups. One group, which is quite clear and distinct from the others, consists of strains of *Zygosaccharomyces* spp., isolated from different species of the genus *Scaptotrigona*, suggesting that these strains belong to the same yeast species previously described for *S. depilis*. A second group consists of *Zygosaccharomyces* strains derived from the brood cells of several bee species of the genus *Melipona*, as well as the species *T. clavipes* and *P. helleri*. In contrast to *Scaptotrigona* isolates, *Zygosaccharomyces* spp. isolated from *Melipona* species probably belong to different species. In turn, *Zygosaccharomyces* isolates from different colonies of the species *T. clavipes* appear to belong to the same species, since they are closely related. Altogether, our findings suggest a widespread symbiotic relationship between species of *Zygosaccharomyces* and SBs, and expand the current knowledge about the classical and non-classical mutualisms between fungi and insects (14). Meliponini are important pollinators of native plants and agricultural crops, playing important ecosystem services. Declines in these pollinators are caused by several factors, including deforestation and pesticides used in agriculture. Pesticides can directly affect bees (15), and our results raise the hypothesis that fungicides used in agriculture could disrupt the bee-yeast symbiosis, indirectly affecting SBs health. Further studies are needed to evaluate the toxicity of these compounds in this symbiotic system to help define public policies for rational use of pesticides in agricultural crops.

**Figure 2.**
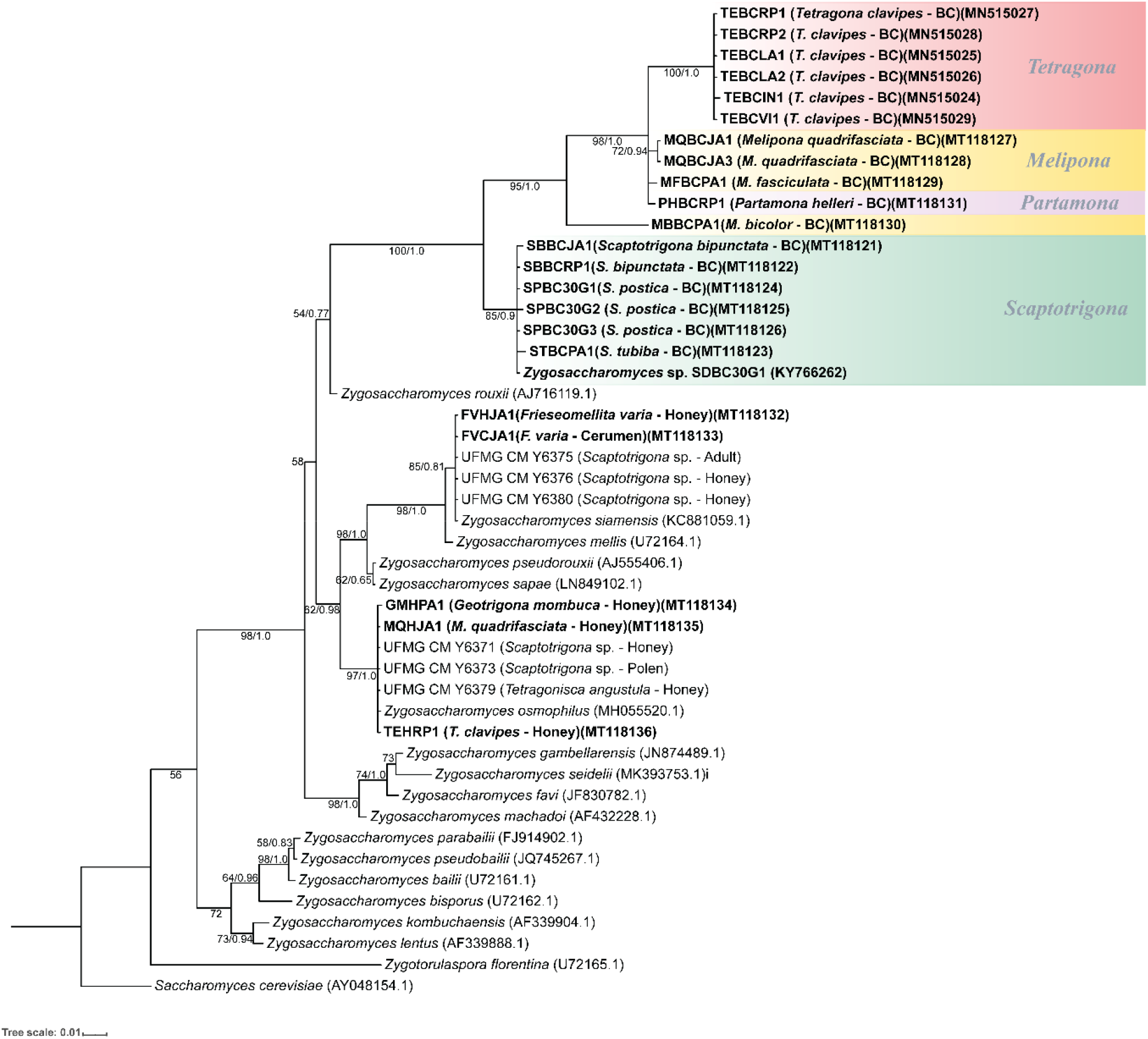
Phylogenetic tree from Bayesian analysis based on sequences of the D1/D2 domain of the 26S rRNA gene of *Zygosaccharomyces* spp. strains isolated from various stingless bee species (highlighted in bold) and from previously described *Zygosaccharomyces* species retrieved from GenBank. Strains isolated from brood cells are highlighted in color, grouped by stingless bee genera. Numbers on branches indicate PP/ML bootstrap values of support for each clade. A total of 540 aligned positions were analyzed. The scale bar represents 0.01 substitutions per nucleotide position. BC: Brood Cell.

## Data availability

The complete data set is presented in the text and the supplemental material.

## Acknowledgments

This research was supported by São Paulo Research Foundation (FAPESP) grants #2013/50954-0 (MTP), #2013/07600-3 (MTP; CEPID-CIBFar), #2018/03650-0 (GTP), #2015/01001-6 (WGPM), by the Conselho Nacional de Pesquisa e Desenvolvimento Tecnológico (CNPq) grants #303792/2018-2 and #307893/2022-7 (MTP); #408733/2021-7 and #406564/2022-1 (CAR), and by Fundação de Amparo à Pesquisa do Estado de Minas Gerais (FAPEMIG) grant APQ-01525-14 (CAR). This study was financed in part by the Coordenação de Aperfeiçoamento de Pessoal de Nível Superior - Brasil (CAPES) - Finance Code 001. Claudia C. de Macedo and Izabel Cristina C. Turatti are acknowledged by their technical support. We also acknowledge the facilities of the Laboratório Multiusuário de Microscopia Confocal – LMMC, FAPESP #2004/08868-0.

## Author Contributions

Conceptualization – MTP, GTP, WGPM, CM, CRP, CAR; Data curation – MTP; Formal analysis – MTP, GTP, WGPM; Funding acquisition – MTP, CAR; Investigation – GTP, WGPM; Methodology – GTP, WGPM, IC, CM; Project administration – MTP; Resources – MTP; Supervision - MTP; Validation – GTP, WGPM; Visualization – MTP, WGPM; Writing - original draft – MTP, GTP, WGPM; Writing - review & editing – MTP, GTP, WGPM, CM, CRP, CAR.

We declare no conflict of interest.

