## Supplemental Information for "Further evidences of an emerging stingless bee-yeast symbiosis"

### Materials and Methods

#### Insect collection and fungal isolation

Permits for collection of biological samples and research were issued by the Brazilian government (SISBIO 46555-6, CNPq 010936/2014-9). Samples of brood cells, honey and cerumen from the nests of different stingless bee species were collected from May 2017 to November 2019 at different cities in Brazil. The GPS coordinates, the information about the isolated microorganisms and the stingless bee species can be found in Table S1.

The growth of white microorganisms in brood cells was visualized in stereomicroscope, and the pseudohyphae were carefully collected with a pointed spatula and aseptically seeded on Petri dishes containing 30 G medium (30 g glucose, 3 g malt extract, 3 g yeast extract and 2 g agar on 100 mL deionized water, pH 6.0) (1). Other parts of the colonies (honey, cerumen, and larval food) were also assessed for fungal isolation. One gram of each material was transferred to an Eppendorf tube containing 200 µL of the 30 G liquid medium (30 g glucose, 3 g malt extract, 3 g yeast extract, and 100 mL deionized water, pH 6.0). Samples were shaken for 30 seconds and transferred to Petri dishes containing 30 G agar medium. Petri dishes were incubated at 29 °C until the beginning of microbial growth and serial cultures were established to obtain pure cultures. Stocks were stored in cryovials at -80 °C in a homogenized mixture of 30 G liquid medium supplied with 20% glycerol.

#### Cell morphology

Images of a small portion of the microorganisms grown in a Petri dish and images directly obtained from the brood cells were registered by stereo microscope (Leica EZ4 W) integrated

5-megapixel cameras to analyze the phenotype of each strain. To analyze cell morphology, cells were harvested using a platinum loop and transferred to a slide containing 50 μL of saline (0.85% NaCl). The cells were mixed on the slide attached to this homogenate and then observed under an optical microscope (Leica ICC50 HD) at 100× magnification.

**Analysis of steroids by fluorescence microscopy**

Staining of cytoplasmic lipid-accumulating organelles, also known as adiposomes, was performed according to the methodology described by Hoiczyk and co-workers (2) and the procedure previously standardized in our laboratory (1). Cells fixed on glass slides at 40 ºC were stained with 50 μL of Nile Red dye (Nile Red, Sigma-Aldrich) for 30 min and dissolved in ethanol (10 μg/mL) in the dark. After this time, cells were quickly washed to remove excess of dye and observed under a fluorescence microscope. Analyzes were performed using the Leica TCS SP5 confocal microscope.

**DNA extraction and fungal identification**

Genomic DNA was extracted following an adapted protocol of Sharma and Singh (3). Fungi were grown in 30 G agar medium at 29 °C until sufficient biomass was achieved. After growth, a portion of each culture was removed and transferred to 2 mL Eppendorf tubes containing 500 μL TE buffer 0.01X. 3 μL RNase was added to digest the RNA, and then 400 μL of solution I (1% sarkosyl, 0.5 M NaCl, 1% sodium dodecyl sulfate (SDS)) also was added. The tubes were kept at 37 °C for 10 min with constant stirring. Then 400 μL phenol:chloroform:isoamyl alcohol 25:24:1 (PCI) was added and mixed by inverting, and centrifuged at 10,000 g for 5 min at 37 °C, followed by careful transfer of the supernatant to a new 1.5 mL Eppendorf tube. Then, 10% of the total volume of sodium acetate (3 M, pH 5.2) and 60% isopropanol were added and mixed gently by inverting the tube 5 to 10 times. Then the tubes were centrifuged at 10,000 g at 37 °C for 5 minutes. The DNA was precipitated in the pellet and the liquid was removed. The pellet was washed with 1 mL of 70% ethanol and centrifuged at 10,000 g for 3 min at 37 °C. The supernatant was removed, and the DNA was resuspended in 20 μL TE 0.01X after drying.

PCR amplification was performed with the 18S SSU rRNA using primers NS1 (5'-GTAGTCATATGCTTGTCTC-3') and NS4 (5'-CTTCCGTCAATTCCTTTAAG-3') (4), 26S LSU rRNA (D1/D2 domains) using primers NL1 (5'-GCATATCAATAAGCGGAGGAAAAG-3') and NL4 (5'-GGTCCGTGTTTCAAGACGG-3') (5), and the ITS (Internal Transcribed Spacer) region containing primers ITS5 (5'-TCCGTAGGTGAACCTGCGG-3') and ITS4 (5'-TCCTCCGCTTATTGATATGC-3') (4). The final reaction volume of 15 μL contained: 8 μL PCRBIO HS Taq Mix (PCRBiosystems), 0.5 μL of each primer, 0.5 μL DMSO, 4.5 μL deionized $H_2O$, and 1 μL DNA (10 ng/ μL). Amplification products were subjected to agarose gel electrophoresis at 1% in 1X TBE buffer and stained with GelRed® (Biotium, Fremont, USA). Purification was performed using the GenElute PCR Clean-up Kit (Sigma-Aldrich).

The sequencing reaction was performed using 2.0 μL 5X buffer, 0.32 μL primer (10 μM), 0.3 μL BigDye® 3.1 (Applied Biosystems, Waltham, USA), approximately 20 ng of purified DNA, and deionized $H_2O$ to achieve a total volume of 10 μL. The thermocycler program consisted of 28 cycles at 95 °C for 15 s, followed by 50 °C for 45 s and 60 °C for 4 min, which were completed and maintained at 10 °C. The sequencing reaction was purified according to the instructions in the BigDye® manual (Applied Biosystems, Waltham, USA).

Sequencing was performed using the 3500 Genetic Analyzer from (Applied Biosystems, Waltham, USA). The sequences were edited in BioEdit 7.2.5 (6) and used to assemble the contigs. The contigs were used to search for homologous sequences in the National Center for Biotechnology Information (NCBI) - GenBank (https://blast.ncbi.nlm.nih.gov/Blast.cgi) database.

**Phylogenetic analyses**

A phylogenetic analysis of the 26S LSU rRNA and a combined phylogeny of 18S and 26S were performed to include the isolated *Zygosaccharomyces* strains and other species previously described to cover most of the genus. After assembly of the contigs, sequences were manually aligned and trimmed in MEGA version X (7). Phylogenetic and molecular evolutionary analyses were reconstructed using Bayesian Inference (BI) in MrBAYES v.3.2.2. (8) with the substitution model general time-reversible with Gamma distribution for rate variation (GTR + I + G) for 26S phylogeny and GTR + G for combined phylogeny. Two separate runs were performed, each consisting of a cold chain and three incrementally heated chains and a Markov Chain Monte Carlo (MCMC) with three million generations. One in each 50 generations were sampled to obtain Bayesian posterior probability (PP) values for the clades. Convergence occurred when the standard deviation (SD) of split frequencies fell below 0.01, and the first 10% of generations in the MCMC samples were discarded as burn-in. The final phylogenetic trees were visualized using iTOL v5 (9).

A maximum likelihood analysis was also conducted on the 26S LSU and combined dataset using the GTR + I + G model. Runs were performed using PAUP 4.0

(https://paup.phylosolutions.com/) software and probability bootstrap support values were generated from 1000 replicates.

The analysis of 26S LSU and the combination of 18S and 26S included 45 and 39 nucleotide sequences, respectively. The first phylogenetic tree was rooted by *Saccharomyces cerevisiae* NRRL Y-12632 and the second one by *Wickerhamomyces anomalus* NRRL Y-366. The sequences are deposited in GenBank, and the deposit numbers are listed in Table 1.

**Ergosterol extraction**

Microorganisms were grown in small Petri dishes (60 mm) containing 30G6 agar medium for 15 days at 29 ºC. Cells were harvested with a loop and transferred to a 2mL Eppendorf tube (previously weighed) containing 1 mL of sterile saline (0.85% NaCl). This suspension was then vortexed for 10 seconds and centrifuged at 18,000 g for 10 minutes. The supernatant was discarded, and the washing procedure was repeated twice. After the last cell wash centrifugation, the supernatant was discarded, and the Eppendorf tube containing the cells was exposed to speed vacuum (Savant, Thermo Scientific) until the cells were completely dried. Each tube containing the samples was weighed and the weight of the empty tube was subtracted to obtain the final mass of each microorganism. For extraction, 250 µL of sterile saline was added to each Eppendorf tube containing the standardized mass. This suspension was vortexed for 20 seconds, then 750 µl of chloroform was added. This mixture was vortexed for 20 seconds and then shaken at 1,500 rpm for 1 hour at room temperature (Agimaxx, Thermo Shaker). The organic phase was collected and made up to a final volume of 1 mL with chloroform. The internal standard (β-sitosterol - 25 µg/mL) was then added and analyzed by gas chromatography coupled to mass spectrometry (GC-MS). Analyzes were performed at NPPNS-FCFRP-USP using a Shimadzu QP -2010 Plus gas chromatograph. A Phenomenex ZB-5MS (ZEBRON) column (30 m x 0.25 mm x 0.25 µm) with a flow rate of 1.50 mL/min, injection temperature of 250 °C, interface of 300 °C, and ionization source of 250 °C in splitless mode was used. The temperature gradient used was 70 °C for 4 minutes and then increased by 10 °C/min to 300 °C, remaining at 300 °C for 15 minutes.

**Extended Results and Discussion**

Usually, fungal filaments are readily visible in the brood cells and occur in the early larval stages. In *Scaptotrigona* spp. and *M. bicolor*, *Zygosaccharomyces* strains grow as long filaments and in substantial amounts, which facilitates isolation. The yeasts associated with *M. fasciculata*, *M. quadrifasciata*, and *P. helleri* form small filaments in small quantities in the first two bee species and in massive quantities in the last one. In contrast to the others, *Zygosaccharomyces* sp. from *T. clavipes* is found in clumps and also in large amounts (Figure 2). The visibility of fungal filaments in brood cells of stingless bees may be limited by the arrangement and size of the brood cell diameter (Figure S1). The strains that originated from different sites in the colonies (honey, pollen, refuse, and adult bees) grew faster during the treatment (at least two days) than the strains that originated from brood cells (at least seven days), suggesting that there are physiological differences between them.

*Zygosaccharomyces osmophilus* was isolated from the honey of *Tetragona clavipes*, *Geotrigona mombuca* and *M. quadrifasciata*, while *Z. siamensis* was isolated from honey and cerumen of the stingless bee *F. varia*.

Since it was not possible to visualize fungal filaments using a stereomicroscope, the cerumen was scraped off with a pointed spatula and the brood cells were washed with 30 G liquid medium for microbial isolation. This allowed the isolation of other fungal genera such as *Leiothecium, Xerochrysium* and *Monascus*. Filamentous fungi such as *Monascus* sp., *M. ruber*, *Xerochrysium* sp. and *Leiothecium* sp. were found in colonies of *Scaptotrigona bipunctata*, *Tetragonisca angustula*, *Friesella schrottkyi*, *Nannotrigona testaceicornis* and *Melipona quadrifasciata*. *Monascus ruber* (SBBCRP2) was isolated from the cerumen of the brood cell of *Scaptotrigona bipunctata*. The other isolates of this genus showed the lowest query coverage and identity percentages with species from the Genbank database and may represent new species.

No fungal strain was isolated from the bees *Frieseomelitta silvestrii*, *F. doederleini*, *Leurotrigona muelleri*, *Melipona quinquefasciata*, and *Oxytrigona tataira*.

*Zygosaccharomyces* species have often been isolated from various parts of bee nests. In honeybees (*Apis mellifera*), different species of *Zygosaccharomyces* have been isolated from bee bread and honey (*Z. mellis*, *Z. rouxii*, *Z. siamensis*, *Z. favi*, and *Z. gambellarensis*) (10, 11), from pollen stored under warm temperatures (12), and as part of the gut microbiota of these bees (13).

*Zygosaccharomyces* is also present in various parts of stingless bee nests. Echeverrigaray and coworkers (14) reported the presence of several species of *Zygosaccharomyces* in honey of stingless bees, such as *Z. mellis* (*M. quadrifasciata*, *Plebeia emerina*, *S. depilis*, and *Tetragonisca fiebrigi*), *Z. rouxii* (*P. droryana*, *P. emerina*, and *T. fiebrigi*), and *Z. siamensis* (*T. fiebrigi*). *Z. siamensis* has also been isolated from the honey of *M. interrupta* (15). Haag and collaborators (16) reported the presence of *Zygosaccharomyces* in the associated gut microbiota of *M. quadrifasciata* through a longitudinal metabarcoding study. Another species, *Zygosaccharomyces machadoi*, was isolated from garbage pellets of the stingless bee *Tetragonisca angustula* (17). Recently, a new species of *Zygosaccharomyces*, described as *Z. osmophilus*, was isolated from unripe and ripe honey and pollen of *S. bipunctata* and from ripe honey of *T. angustula* (18). Similarly, *Zygosaccharomyces* species were also isolated from honey and cerumen of stingless bees in the present study. *Zygosaccharomyces osmophilus* was isolated from honey of *Tetragona clavipes*, *Geotrigona mombuca*, and *M. quadrifasciata*, and *Z. siamensis* from honey and cerumen of stingless bee *F. varia*. Thus, there are different species of *Zygosaccharomyces* associated with stingless bees and present in different locations in the nests.

In addition to what was discussed in the Observation session, it was found that the isolation of *Zygosaccharomyces* sp. from different colonies of the same stingless bee species, as in *T. clavipes*, is further evidence that this microorganism is indeed present in this system and may have a specific function. The branch consisting only of isolates of *Zygosaccharomyces* spp. from brood cells of different stingless bee species is present in both phylogenetic trees (Figure 2 and Figure S4). These lineages are closely related, sharing a common ancestor, but included in a separate branch from other previously described *Zygosaccharomyces*.

*Zygosaccharomyces* sp. SDBC30G1 was compared with *Z. rouxii* (the closest described species to the isolated brood cell branch), and a difference of 49 substitutions (8.3%) was observed in a dataset of 540 nucleotides (nt) (Figure S5). In addition, the sequences of *Zygosaccharomyces* isolates from the brood cell of the other SBs species were compared with *Zygosaccharomyces* sp. SDBC30G1. The nt sequences of the SBs isolates from the genus *Scaptotrigona* showed an average percent identity of 99.5% reinforcing that they are the same species as *Zygosaccharomyces* sp. SDBC30G1. The sequences of isolates from the genus *Melipona* and the species *P. heleri* showed an average difference of 62 (11.5%) substitutions in a dataset of 540 nt. The same occurs with the *T. clavipes* isolates, which show an average difference of 76 (14.2%) substitutions in a dataset of 540 nt. These data support that we have candidates for new and distinct species of *Zygosaccharomyces* in the branch of brood cell isolates from different species of SBs.

Our results suggest that *Zygosaccharomyces* species found in brood cells of stingless bees are specific to this system, despite differences in host insect morphology, behavior, and nest structure. These yeasts, although derived from different bee species, are likely associated with a specific nutritional function for bee larvae, as previously demonstrated for *S. depilis* (1). Therefore, these yeasts may have been evolutionarily selected to fulfill this nutritional function for these host bees. *Zygosaccharomyces* spp. in turn find a favorable environment for their growth in the brood cell.

Other isolation techniques were used to isolate the microorganisms, such as scraping the cerumen from the brood cell or washing the entire brood cell with saline, mainly in those species of SBs that did not have visible fungal filaments. This allowed the isolation of other species of non-*Zygosaccharomyces* fungi. Due to the use of this non-specific technique, it is not possible to conclude that these different isolated fungi have the same biological function as *Zygosaccharomyces* species for larval development of stingless bees.

*Monascus* sp. SBBCRP2 isolated from the brood cell of *S. bipunctata* is morphologically similar to the strain of *Monascus ruber* SDBC1 previously isolated from the cerumen of the brood cell of *S. depilis*. *M. ruber* SDBC1 was found to negatively modulate *Zygosaccharomyces* sp. growth by producing lovastatin. Moreover, *M. ruber* SDBC1 exerts a similar function on *Candida* sp., which is also present in this system, by producing monascin (19). *Monascus* strains have been previously isolated from honey, pollen, and inside the nest of *M. scutellaris*, including *Monascus recisfensis* (pollen), *M. ruber* (in the nest), *M. mellicola* (honey, pollen, and in the nest), and *M. flavipigmentosus* (pollen and in the nest) (20) indicating a possible ecological interaction of this genus with stingless bees.

**Table S1.** Data from strains of microorganisms isolated from different stingless bees nest sites.

| Stingless Bee Species | Sample | Sample ID | GenBank Access Number (18S) | GenBank Access Number (26S) | Species | City and State[#] (Brazil) | GPS coordinates |
|---|---|---|---|---|---|---|---|
| ***Scaptotrigona bipunctata**** | Brood cell | SBBCJA1 | MT118313 | MT118121 | *Zygosaccharomyces* sp. | Jaguariúna-SP | 22.732714 S 47.018359 W |
| | | SBBCRP1 | MT118312 | MT118122 | *Zygosaccharomyces* sp. | Ribeirão Preto-SP | 21.169779 S 47.859670 W |
| | | SBBCRP2 | - | - | *Monascus ruber* | Ribeirão Preto-SP | 21.169779 S 47.859670 W |
| ***Scaptotrigona postica**** | Brood cell | SPBC30G1 | MT118315 | MT118124 | *Zygosaccharomyces* sp. | Belém-PA | 1.436129 S 48.449181 W |
| | | SPBC30G2 | MT118316 | MT118125 | *Zygosaccharomyces* sp. | Belém-PA | 1.436129 S 48.449181 W |
| | | SPBC30G3 | MT118317 | MT118126 | *Zygosaccharomyces* sp. | Belém-PA | 1.436129 S 48.449181 W |
| ***Scaptotrigona tubiba**** | Brood cell | STBCPA1 | MT118314 | MT118123 | *Zygosaccharomyces* sp. | Passos-MG | 20.720259 S 46.613014 W |
| ***Tetragonisca angustula*** | Brood cell | TABCJA1 | - | - | *Xerochrysium* sp. | Jaguariúna-SP | 22.732714 S 47.018359 W |
| ***Tetragona clavipes**** | Honey | TEHRP1 | MT118327 | MT118136 | *Zygosaccharomyces mellis* | Ribeirão Preto-SP | 21.169779 S 47.859670 W |
| | Brood cell | TEBCRP1 | MT118309 | MN515027 | *Zygosaccharomyces* sp. | Ribeirão Preto-SP | 21.169779 S 47.859670 W |
| | | TEBCRP2 | MT118310 | MN515028 | *Zygosaccharomyces* sp. | Ribeirão Preto-SP | 21.169779 S 47.859670 W |
| | | TEBCLA1 | MT118307 | MN515025 | *Zygosaccharomyces* sp. | Luís Antonio-SP | 21.575217 S 47.746817 W |
| | | TEBCLA2 | MT118308 | MN515026 | *Zygosaccharomyces* sp. | Luís Antonio-SP | 21.575217 S 47.746817 W |
| | | TEBCIN1 | MT118306 | MN515024 | *Zygosaccharomyces* sp. | Indaiatuba-SP | 23.066368 S 47.170355 W |
| | | TEBCVI1 | MT118311 | MN515029 | *Zygosaccharomyces* sp. | Viçosa-MG | 20.758624 S 42.868174 W |
| ***Frieseomelitta varia*** | Honey | FVHJA1 | MT118324 | MT118132 | *Zygosaccharomyces siamensis* | Jaguariúna-SP | 22.732714 S 47.018359 W |
| | Cerumen | FVCJA1 | MT118325 | MT118133 | *Zygosaccharomyces siamensis* | Jaguariúna-SP | 22.732714 S 47.018359 W |

| | | | | | | | |
|---|---|---|---|---|---|---|---|
| | Brood cell | FVBCRP1 | - | - | - | Ribeirão Preto-SP | 21.169779 S 47.859670 W |
| | | FVBCRP2 | - | - | - | Ribeirão Preto-SP | 21.169779 S 47.859670 W |
| ***Frieseomelitta silvestrii*** | Brood cell | - | - | - | - | Ribeirão Preto-SP | 21.169779 S 47.859670 W |
| ***Frieseomelitta doederleini*** | Brood cell | FDBCPA1 | - | - | - | Passos-MG | 20.720259 S 46.613014 W |
| ***Friesella schrottkyi*** | Brood cell | FSBCJA1 | - | - | *Leiothecium* sp. | Jaguariúna-SP | 22.732714 S 47.018359 W |
| | | FSBCJA2 | - | - | *Monascus* sp. | Jaguariúna-SP | 22.732714 S 47.018359 W |
| ***Geotrigona mombuca*** | Honey | GMHPA1 | MT118326 | MT118134 | *Zygosaccharomyces mellis* | Passos-MG | 20.720259 S 46.613014 W |
| ***Leurotrigona muelleri*** | Brood cell | - | - | - | - | Jaguariúna-SP | 22.732714 S 47.018359 W |
| ***Nannotrigona testaceicornis*** | Brood cell | NTBCJA1 | - | - | *Monascus* sp. | Jaguariúna-SP | 22.732714 S 47.018359 W |
| | | NTBCJA2 | - | - | *Monascus* sp. | Jaguariúna-SP | 22.732714 S 47.018359 W |
| | | NTBCRP1 | - | - | *Monascus* sp. | Ribeirão Preto-SP | 21.169779 S 47.859670 W |
| ***Plebeia droryana*** | Brood cell | PDBCJA1 | - | - | *Monascus* sp. | Jaguariúna-SP | 22.732714 S 47.018359 W |
| ***Melipona quadrifasciata**** | Honey | MQHJA1 | - | MT118135 | *Zygosaccharomyces mellis* | Jaguariúna-SP | 22.732714 S 47.018359 W |
| | Brood cell | MQBCJA1 | MT118318 | MT118127 | *Zygosaccharomyces* sp. | Jaguariúna-SP | 22.732714 S 47.018359 W |
| | | MQBCJA2 | - | - | *Monascus* sp. | Jaguariúna-SP | 22.732714 S 47.018359 W |
| | | MQBCJA3 | MT118319 | MT118128 | *Zygosaccharomyces* sp. | Jaguariúna-SP | 22.732714 S 47.018359 W |
| | | MQBCPA1 | - | - | *Zygosaccharomyces* sp. | Passos-MG | 20.720259 S 46.613014 W |
| ***Melipona quinquefasciata*** | Brood cell | - | - | - | - | Passos-MG | 20.720259 S 46.613014 W |
| ***Melipona fasciculata**** | Brood cell | MFBCPA1 | MT118321 | MT118129 | *Zygosaccharomyces* sp. | Passos-MG | 20.720259 S 46.613014 W |
| ***Melipona bicolor**** | Brood cell | MBBCPA1 | MT118322 | MT118130 | *Zygosaccharomyces* sp. | Passos-MG | 20.720259 S 46.613014 W |
| ***Oxytrigona tataira*** | Brood cell | - | - | - | - | Passos-MG | 20.720259 S 46.613014 W |
| ***Partamona helleri**** | Brood cell | PHBCRP1 | MT118323 | MT118131 | *Zygosaccharomyces* sp. | Ribeirão Preto-SP | 21.169779 S 47.859670 W |

* Fungal filaments were observed in the brood cell; #States in Brazil: MG - Minas Gerais; PA – Pará; SP - São Paulo.

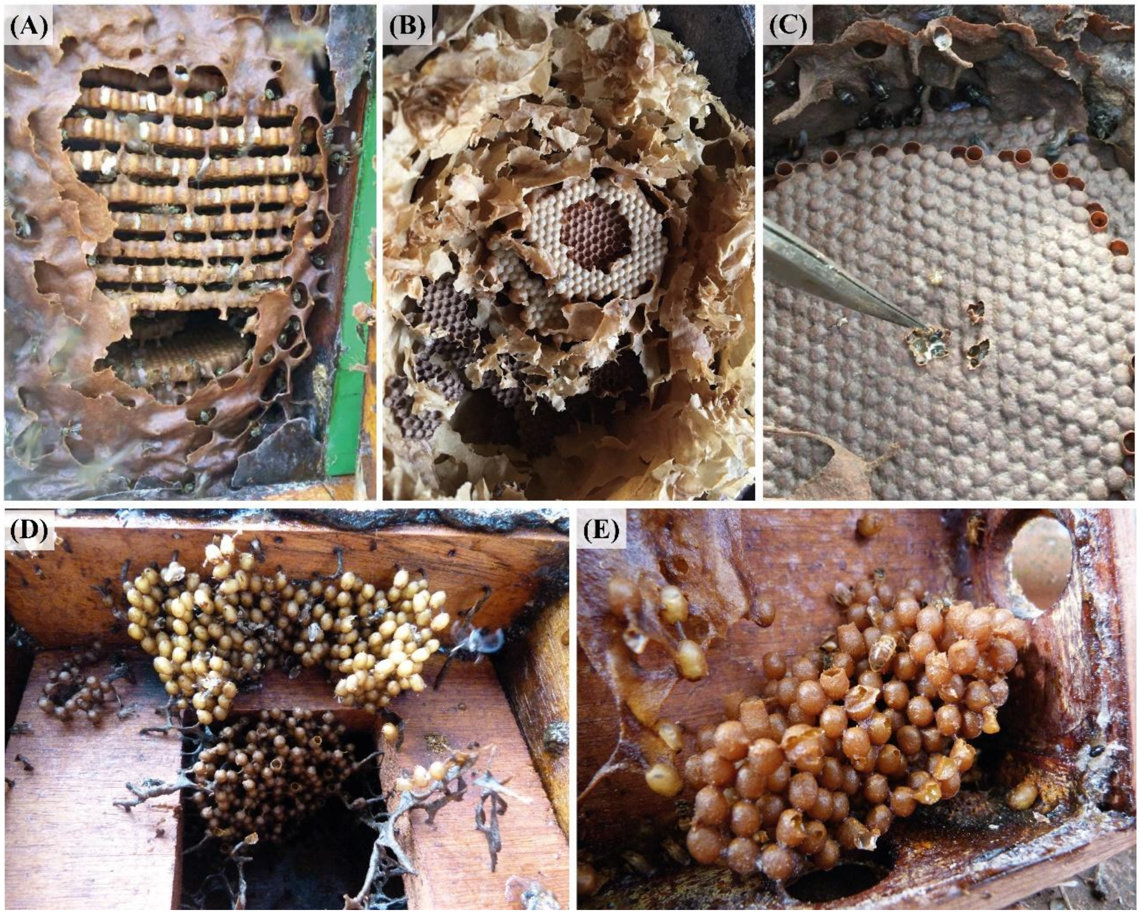


**Figure S1.** Brood cells of different stingless bee species. (A) *Tetragona clavipes* (B) *Nanotrigona testaceicornis* (C) *Scaptotrigona bipunctata* (D) *Frieseomelitta silvestrii* (E) *Friesemelitta varia*. The brood cells of A, B, and C are arranged in the form of disks, while D and E are arranged in the form of clusters.

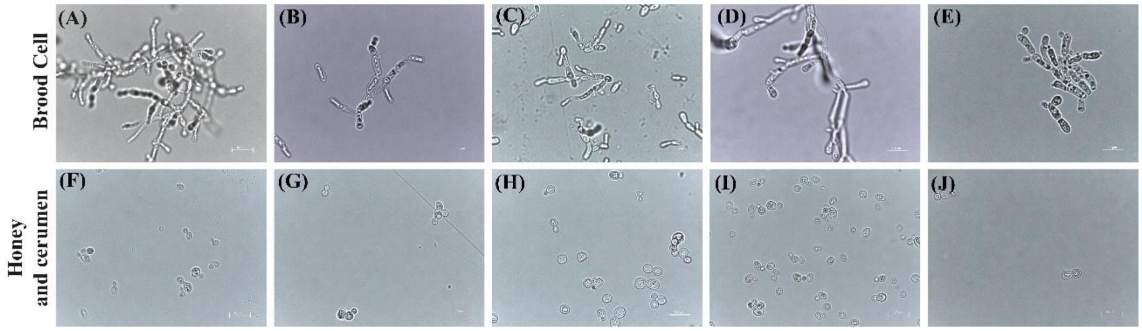


**Figure S2.** Comparative analysis of cells of *Zygosaccharomyces* spp. by optical microscopy at 100x magnification. Images (A), (B), (C), (D) and (E) obtained from strains isolated from brood cells (A: *M. bicolor*; B: *T. clavipes*; C: *P. helleri*; D: *S. bipunctata* and E: *S. tubiba*). Strains (F-I) were isolated from other sources: (F) from cerumen of *F. varia,* and (G), (H), (I) and (J) from honey (G: *F. varia*, H: *G. mombuca*, I: *T. clavipes* and J: *M. quadrifasciata*). The strains from the brood cells show pseudohyphae formation while strains from other parts of the colony present ovoid cells.

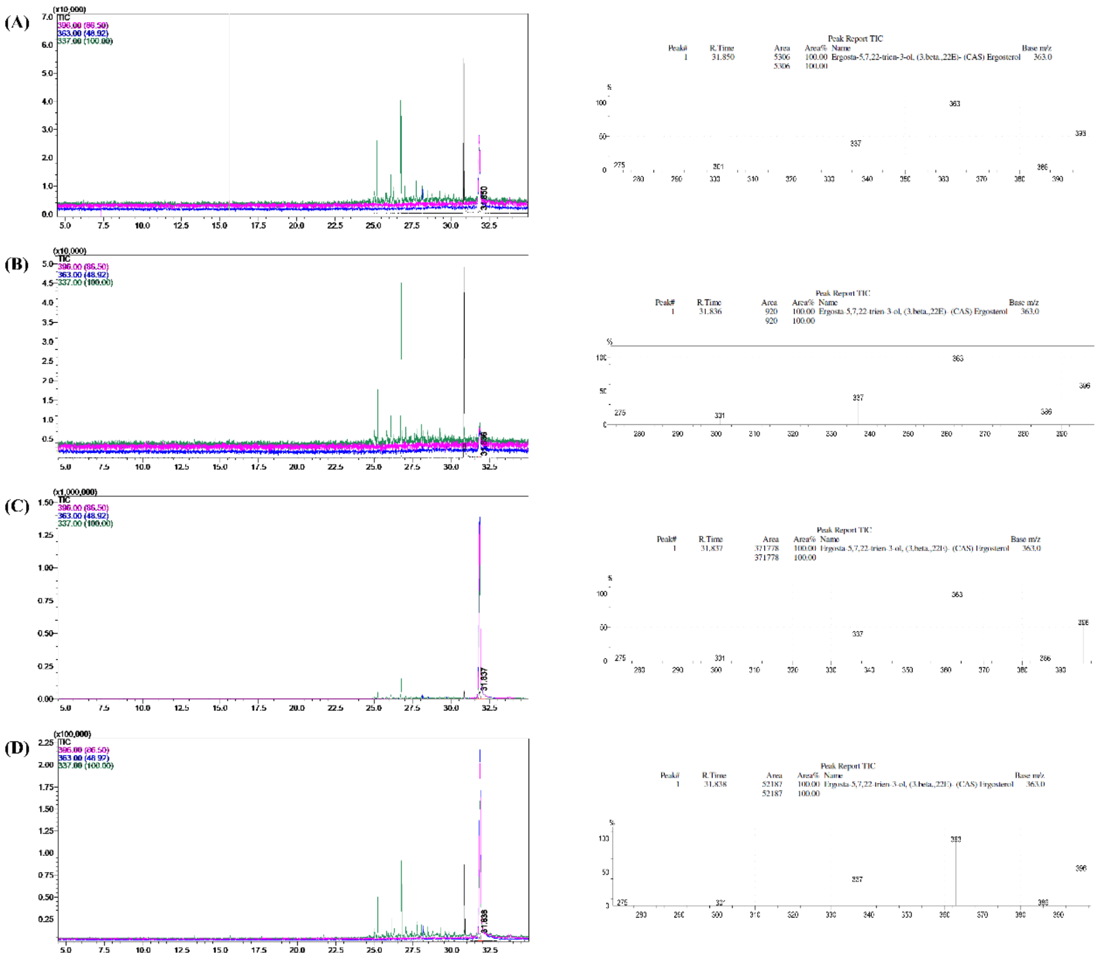


**Figure S3.** Ergosterol analysis by GC-MS of *Zygosaccharomyces* spp. strains from different stingless bee species: (A) *Melipona bicolor,* (B) *Scaptotrigona bicolor,* (C) *Melipona quadrifasciata,* (D) *Tetragona clavipes*. The images contain the chromatogram with the ergosterol retention time highlighted, followed by the identification suggested by the program and the ergosterol equivalent fragment ions.

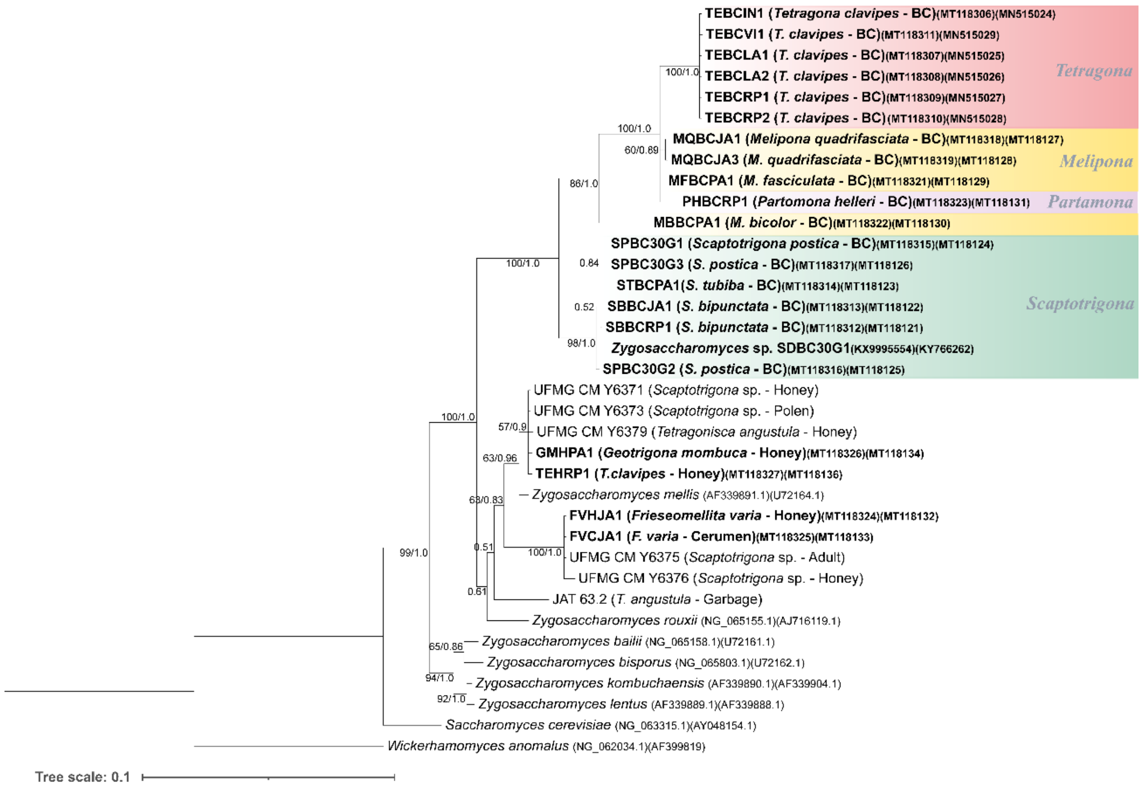


**Figure S4.** Bayesian inference based on concatenated sequences of the 18S gene and D1/D2 domain of the 26S rRNA gene from *Zygosaccharomyces* spp. strains isolated from stingless bee species (highlighted in bold) and previously described *Zygosaccharomyces* species retrieved from GenBank. Strains isolated from brood cells are highlighted in color, grouped by stingless bee genera. Numbers on branches indicate PP/ML bootstrap values of support for each clade. A total of 1640 aligned positions were analyzed. The scale bar represents 0.1 substitutions per nucleotide position.

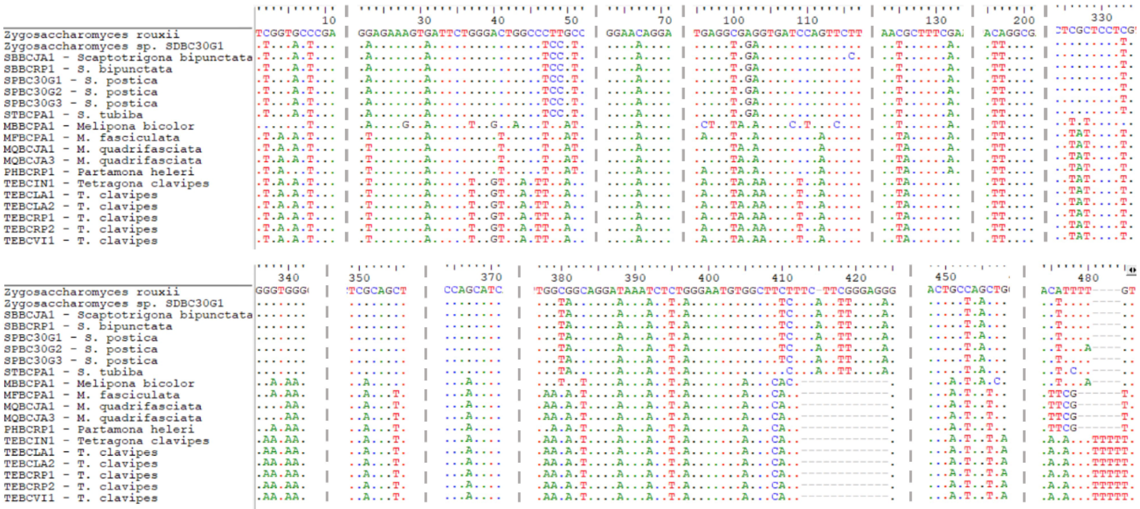


**Figure S5.** Comparison of the 26S gene nucleotide sequences of the brood cell isolates of different SBs species, *Zygosaccharomyces* sp. SDBC30G1 and *Z. rouxii*. Sequences were aligned using MEGA version X (7) and visualized in BioEdit 7.2.5 (6).
